# Automatic Consistency Assurance for Literature-based Gene Ontology Annotation

**DOI:** 10.1101/2021.05.26.445910

**Authors:** Jiyu Chen, Nicholas Geard, Justin Zobel, Karin Verspoor

## Abstract

**Background:** Literature-based gene ontology (GO) annotation is a process where expert curators use uniform expressions to describe gene functions reported in research papers, creating computable representations of information about biological systems. Manual assurance of consistency between GO annotations and the associated evidence texts identified by expert curators is reliable but time-consuming, and is infeasible in the context of rapidly growing biological literature. A key challenge is maintaining consistency of existing GO annotations as new studies are published and the GO vocabulary is updated.

**Method:** In this work, we introduce a formalisation of biological database annotation inconsistencies, identifying four distinct types of inconsistency. We propose a novel and efficient method using state-of-the-art text mining models to automatically distinguish between consistent GO annotation and the different types of inconsistent GO annotation. We evaluate this method using a synthetic dataset generated by directed manipulation of instances in an existing corpus, BC4GO.

**Results and Conclusion:** Two models built using our method for distinct annotation consistency identification tasks achieved high precision and were robust to updates in the GO vocabulary. We provide detailed error analysis for demonstrating that the method achieves high precision on more confident predictions. Our approach demonstrates clear value for human-in-the-loop curation scenarios.

**Data availability:** The synthetic dataset, and the code for generating it are available at https://github.com/jiyuc/BioConsistency.

## Introduction

The Gene Ontology (GO) is a framework used to uniformly describe gene function and support the representation of biological systems, based on a set of hierarchical controlled vocabularies [1, 2]. GO annotation of genes involves two major components: the *GO information*, which includes GO terms and their definitions or descriptions, and *supportive evidence*, which includes the coding regions on a genomic sequence, or a reference to a document describing experimental findings relating to gene product function. Sequence-based GO annotations are produced by comparing different sequences drawn from coding regions; this process can transfer GO terms from an old sequence to a new because two genomic sequences with structurally closer coding regions tend to produce gene products with similar functions [3, 4]. Literature-based GO annotations (GOA) are produced by reviewing the description of experiments in research papers, selecting appropriate GO terms for the experimental findings, and labelling the annotation with a GO evidence code^[1]^ [5–8] indicating the nature of the evidence. While the majority of sequence-based and literature-based GO annotations are automatically produced, the most reliable are manually annotated by expert curators. There is a pressing need of reliable tools for automatic curation of GOA as the volume of biological data continues to go.

There are currently around eight million GOA across 4, 743 species recorded in the GO Consortium Database^[2]^. However, fewer than 2% of these are manually curated; these are linked to 162, 459 publications [9]. Automatic GO curation is efficient but the existing benchmarks are unreliable [9–11]. Furthermore, annotation tools that target a fixed set of terms cannot satisfy the open-world assumption, which requires that the collection of GO terms be updated with the discovery of new gene functions [11]. For example, the GO categoriser (GOCat) is based on a closed world assumption, which is that all relevant terms have been previously observed. It relies on a K-Nearest Neighbour algorithm to compare the semantic similarity of an existing GO annotated evidence text and a new text describing a gene function [10]. GOCat uses several strategies to select GO terms from the old evidence text and rank by relevancy to annotate the new text. However, if the new text describes a gene function that has large semantic distance from any existing evidence text, GOCat will fail to shortlist GO terms and thus will skip this functional annotation. Also, tools based on the closed-world assumption may be biased towards frequently selected GO terms [11, 12], such as assigning “protein binding (GO:0005515)” to a large proportion of genes [13]. Another tool called ConceptMapper [14] utilises dictionary-based concept recognition to achieve competitive performance in annotating GO concepts on the Colorado Richly Annotated Full Text (CRAFT) corpus [15, 16]. However, this tool cannot recognise GO concepts that do not explicitly occur as phrases within evidence texts. For example, “positive regulation of vesicle fusion (GO:0031340)” cannot be recognised from “Rat SYT1 gave rise to efficient Ca2+-promoted fusion activity”.

GO annotation is not a one-time process [11]. After a GO term has been assigned to a gene product and linked to evidence, database curators need to continue to monitor the consistency of this annotation against new findings. For example, if a gene product was previously published as having negatively regulated behaviour but is later reported as being uncertain, then this GO annotation should be removed from the database. Poor-quality records within databases can cause cascading errors [17] that in turn may lead to significant negative impact to many down-stream tasks such as gene expression analysis. However, existing studies largely focus on methods for efficient GO annotation enrichment, with less emphasis on maintaining the quality of annotations that have already been recorded within databases.

There are many challenges to assessing the quality of GOA [18]. Some researchers have estimated a relatively high error rate of GO-curated sequence annotations [19]. A series of quality issues in literature-based GO annotations have also been identified. Some GOA are reported as being assigned to unsupportive evidence texts [20]. Some curators find difficulties in selecting informative GO terms at the proper level of specificity [9]. These quality issues can be seen as reflecting inconsistencies b tween GO information and evidence information. No tools have been proposed for automatic evaluation of the correctness of evidence code selection. The current solution is to create comprehensive curation guidelines and manually ensure annotation consistency. However, this is unscaleable [7, 21, 22].

To improve the efficiency and quality of GO annotation, we propose a novel method that uses text mining for the maintenance of literature-based GOA consistency. The method can automatically distinguish consistent GOA and four major types of inconsistencies as well as satisfying the GO open-world assumption. We model GOA (in)consistency as typed pairwise relationships between GO information/evidence code and associated evidence text. We formalise four primary types of GOA inconsistencies that violate curation guidelines [7, 21] or reported by previous researchers (Table 1): the contradictory description of gene regulation function (Type A); over-specific (or over-informative [9]) selection of gene ontology terms (Type B); inappropriate selection of unsupportive texts as evidence (Type C) [20]; and erroneous selection of experimental typed evidence code (Type D). Types A–C involve inconsistencies related to GO term selection while Type D relates to the broader nature of the evidence for the annotation; we model these groups separately. The formalisation of four types of GOA inconsistencies can help curators address detailed exploratory analysis of database consistency issues.

**Table 1.** Examples of four types of inconsistent Gene Ontology Annotations; 3 term-related and 1 related to evidence codes.

|  |  |
| --- | --- |
| Term Inconsistency | **Type A – Contradictory description of gene regulation function**<br>*Evidence:* As expected, rat <u>SYT1</u> gave rise to efficient Ca<sup>2+</sup>-promoted fusion activity.<br>*GO term:* *negative regulation of vesicle fusion (GO:0031339)*<br>*Inconsistency:* The evidence describes that SYT1 has a positive regulation of vesicle fusion which is contradictory to the negative expression in the selected GO term. |
|  | **Type B – Over-specific GO term selection**<br>*Evidence:* Recent studies from this laboratory have provided strong evidence that endogenous GRK2 and GRK6 can regulate the responsiveness of M1 mACh receptor signalling in cultured rat hippocampal neurons.<br>*GO term:* *negative regulation of G-protein coupled receptor protein signaling pathway (GO:0045744)*<br>*Inconsistency:* The provided evidence only support the regulation of G-protein coupled receptor protein singling pathway while its negativity is unknown. Thus, the annotated GO term is over-specific. |
|  | **Type C – Unsupportive evidence text**<br>*Evidence:* In order to characterise the most prominent protein changes that arise in livers from rats fed control or ethanol-containing diets with or without betaine supplementation, <u>cytosolic</u> liver proteins were resolved by 1D PAGE.<br>*GO term:* *cytosol (GO:0005829)*<br>*Inconsistency:* The evidence text has the mention of a GO concept “cytosol” but does not express the cellular component information of any gene product thus the evidence text does not correctly support the selected GO term. |
| Code inconsistency | **Type D – Erroneous selection of experiment type GO evidence code**<br>*Evidence:* At 22h after pollination, we found pollen tubes in 37.5% of the ovaries following pollinations by <u>EXPB1</u> pollen (Fig 4). In contrast, we found no pollen tubes in the ovaries at 22h after pollination when the silks were pollinated by expb1 pollen. Together these data indicate that the <u>expb1</u> pollen grows more slowly in vivo than the EXPB1 pollen.<br>*Evidence code:* IGI<br>*GO term:* *pollen tube growth (GO:0009860)*<br>*Inconsistency:* The evidence indicate the annotation is based on allelic variation and the experiment is conducted by comparing the single gene to the alleles of the same gene. Thus, the correct evidence code should be IMP instead of IGI. |

To evaluate our method, we generate acollection of GOA instances that fall into each (in)consistency category by directed manipulation of instances in the evidencebased BC4GO corpus [23]. We fine-tune two BioPubMedBERT models [24] to distinguish consistent GOA from the three kinds of term inconsistency (**Model-Term**) and from evidence-code inconsistency (**Model-Code**). We propose a simply strategy to extend BioPubMedBERT with additional layer to encode section information marked by the location of evidence text in the article during fine-tuning. The performance of Model-Term and Model-Code are evaluated using *Precision*, *Recall* and *F*_1_ metrics grounded at each evidence text on a test set that is independently generated from 49 full-text articles. Model-Term achieved 0.69 and Model-Code achieved 0.52 *micro-Precision* overall. We optimise training data using a term-overlap similarity measure and improve the ability to distinguish consistent GOA from other types of inconsistencies. We find a significant improvement in precision among each type of (in)consistency when the uncertainty (Shannon entropy) of predicted outcomes decreases.

To identify the typical linguistic features that are influencing the models’ performances, we undertake error analysis based on a linguistic test suite approach [25, 26] and find the length or typical composition structure of GO terms, the occurrence of digits or Roman characters in the GO term, the length of evidence text versus the length of the GO definition text, and overlaps between a GO term and evidence text may all influence the model’s prediction uncertainty and have overall consequences for model performance. Together these outcomes demonstrate the value of our methods as an organising framework, and for improving the efficiency and accuracy of human-in-the-loop GOA curation.

## Background

To provide context for our methods, we introduce the Gene Ontology and describe the existing evidence-based corpus that we exploit in our method and experiments. We also introduce different methods for measuring the semantic similarity between naturally written texts within documents, or different GO terms, modelled on a directed acyclic graph (DAG); some of these are used in our methods. Finally, we discuss the design of linguistic test suites and measurement of prediction uncertainty for post-hoc error analysis. There are no prior methods for automatic maintenance of literature-based GOA consistency, to the best of our knowledge, but as we discuss there are several relevant resources.

The GO is a controlled vocabulary developed to uniformly describe the molecular activity of a gene product (*molecular function*) in a specific location of cell (*cellular component*) and how it contributes to a broad biological objective (*biological process*) [22]. The GO has a hierarchical structure and is modelled as a DAG with terms as nodes and relations between the terms as edges.^[3]^ Parent terms are broad while child terms express more specific information; for example, “suckling behavior (GO:0001967)” is a child term of “feeding behavior (GO:0007631)”, which indicates a more specific form of food intake via nourishment from the breast. Curators need to make sophisticated inferences in order to select the proper specific level of GO term from the hierarchical graph, balancing the need to specify the gene function as precisely as possible against the risk of exceeding the level supported by the evidence.

The BC4GO corpus was created for the GO annotation task in BioCreative IV [23]. In contrast to a mention-based GO corpus such as CRAFT [15], BC4GO mirrors the real-world GO curation scenario, providing each GO annotation with traceable evidence grounded at sentence level within literature. For example, the GO term “growth (GO:0040007)” commonly appears within articles but not every sentence that mentions “growth” is truly supportive gene function evidence. The CRAFT corpus includes annotations of every appearance of “growth” as a GO concept but these are largely not directly relevant for GO curation. The BC4GO corpus categorises evidence sentences into either experiment type or summary type. The experimenttype sentences describe details of how an experiment was conducted and can be used to produce a complete GO annotation by referring to the GO definition and the decision tree for evidence code selection.^[4]^ The summary-type sentences only describe the results of experiments and are used only to infer the selection of GO terms while the evidence code is labelled with “NONE”. Evidence that spans multiple sentences is extracted and concatenated as a single long sentence.

BioSentVec [27] is a sentence semantics representation model pre-trained on a vast volume of PubMed articles and clinical notes [28]. It can transform naturally written sentences into a lower-dimensional vector representation called sentence embeddings. Their model, utilising vectors of dimension 700, achieved competitive results in several biomedical sentence pair similarity prediction tasks [29, 30]. However, the performance of applying BioSentVec for the consistency estimation of two sentences in different level of biomedical information specificity is unknown.

BioPubMedBERT [24] is a contextual representation benchmark pre-trained on domain-specific full-text PubMed articles. It achieved competitive performance in many relation prediction tasks, such as the extraction of drugdrug interactions [31], gene-disease associations [32], and sentence-pair similarity estimation [29]. It uses special tokens “[CLS]” and“[SEP]” to mark the boundary of an entity pair and predict their relation type by mapping the last layer of “[CLS]” encoding into a linear layer for multi-type relation classification. However, the suitability of applying BioPubMedBERT model for open-world consistency inference of sentence pairs is unknown.

Previous work proposed a linguistic test suite for assessing the performance of automatic ontology concept recognition systems [25, 26]. The test suite is designed to extract a set of linguistic features of ontology terms such as the number of English words in the ontology term, the occurrence of digits or Roman characters, or the length of associated evidence text. These linguistic features may impact the model’s prediction uncertainty, which is a metric broadly used in the active learning field [33]. The uncertainties can be represented by the model’s probability or entropy [34] for each prediction, with a lower probability or higher entropy indicating greater uncertainty. In principle, a robust model should perform best on more certain predictions, while flagging of various degrees of quality warnings to humans in real-world curation setting. The estimates of uncertainty also provides possibilities to quantify the model’s performance during error analysis.

## Method

Our approach combines a specific data source with modelling methods. We now introduce each of these components.

### Data

There are no existing GOA resources with labelling of different types of (in)consistencies grounded at evidence level. Thus, to obtain suitable data, we transform consistent GOA available in the BC4GO corpus, generating instances of the four types of inconsistencies we have identified. We use 100 full-text articles from BC4GO to generate a training set and 49 articles to generate a test set. We randomly sample 20% of the generated instances from the training set to form a development set. By doing so, the generated test set is assured to be independent for post-modelling evaluation. We ensure that over 75% of the selected GO terms in the test set do not occur in the training or development set, enable the evaluation of the GO open-world assumption.

To simplify our study, we focus on detecting the main single type of inconsistency in each individual record. Thus, we assume that a GOA can only be in one type of (in)consistency, and further that they are independent of each other. We assume the gene product (usually represented by unique GeneID) and the organism described by the evidence text are consistent with that annotated by the GO term and only focus on detecting the (in)consistencies in gene function descriptions (Eq 1, where *y*_*_ denotes a specific type of (in)consistency, *ϵ* denotes evidence information, *θ* denotes GO information, and *γ* denotes evidence code).

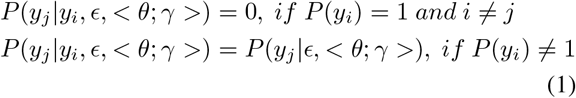

Each GOA instance contains two major components: GO information including the GO term ID, the GO term string and the term definition; and evidence information including evidence texts and spans, evidence codes, section information marked by the location of the evidence text in the article, gene name, gene identification, and gene synonyms. All information is directly extracted from BC4GO annotations or retrieved from the database using QuickGO [35]. The process for preparing each type of (in)consistent instance is as follows:

- ***Consistent* GOA from BC4GO.** We extract GO annotations from full-text articles within BC4GO and transform them to produce instances of consistent GOA. We concatenate evidence text that spans more than one sentence in any GO annotation into a single sentence. We use QuickGO [35] to retrieve any information that was not originally provided in BC4GO annotations such as the GO definition. We remove any GOA in which the GO term is indicated as being obsolete on QuickGO.
- **Type A – *Contradictory* description of gene regulation function.** We apply keyword matching on GO terms in the transformed consistent GOA instances and swap the mention of any “positive regulation” with “negative regulation” and vice versa. We use the manipulated GO term to retrieve associated information such as GO identification, GO definition, and GO synonyms using QuickGO.
- **Type B – *Over-specific* GO term.** We retrieve a list of direct descendants with either “is a” or “part of” relationship to each GO term in consistent GOA instances using QuickGO and manually assure these descendants are over-specific against the evidence texts. We use two alternative strategies to select an overspecific GO term from the retrieved descendants and use it to manipulate the consistent GOA into Type B GOA. 1) We replace the GO term in a consistent GOA with a randomly selected over-specific descendant; 2) We replace the GO term in a consistent GOA with the direct descendant of that term that has the greatest word overlap [36]. For example, “feeding behavior (GO:0007631)” has one overlapping word with descendant “suckling behavior (GO:0001967)” and two overlapping words with descendant “regulation of feeding behavior (GO:0060259)”. Thus, the second descendant will be selected for the replacement.
- **Type C – *Unsupportive* evidence text.** We produce unsupportive variants of each consistent GOA by replacing the evidence sentence with another piece of semantically similar but unsupportive text from the same article. The GO term string in the original consistent GOA occurs as keywords within the chosen text but does not express meaningful gene function information. To find these texts, we refer to the task description of BioCreative IV [20] which states that text that is not annotated with a GO term can be treated as unsupportive. We extract unsupportive texts from the BC4GO corpus by article and segment them into sentences using TextBlob [37]. Each segment is considered as a piece of unsupportive evidence. To find certain pieces of texts that may be confused with valid evidence texts in each consistent GOA, we first apply GO concept recognition as implemented by [16] in the CCP-NLP-Pipelines to recognise any mention of a GO term in these unsupportive evidence sentences. We then pair each GO concept recognised unsupportive sentence with *consistent* GOA instance in the same article that share the same GO concept. In order to select an unsupportive sentence that is most similar to the evidence sentence in the consistent GOA, we represent the sentences as embeddings using BioSentVec [27] and calculate the cosine similarity between each pair of matched evidence sentence and unsupportive sentence. We produce the final instance by replacing the evidence sentence in the consistent GOA with the unsupportive sentence that is most similar. We update information in manipulated GOA such as evidence section information and formalise it into Type C GOA instance.
- **Type D – *Erroneous evidence code***. We select consistent GOA instances where evidence sentences are experiment type, based on an experimental type evidence code label. We exclude the selection of summary-type evidence sentences as they do not support the selection of an evidence code. We replace the evidence code with another code randomly selected from the decision tree mentioned in Section. For example, we replace “IMP” with “IGI” in Table 1 (Type D inconsistency example).

After generating GOA instances, we manually confirmed the true (in)consistency of each automatically generated instance and the associated information. While we did not have a formal manual annotation process, our approach of targeted manipulation of annotated examples leads to reliable labels.

The statistics of the generated dataset is shown in Table 2. The generated dataset, and the code for generating it are available at https://github.com/jiyuc/BioConsistency.

**Table 2.** The number of generated instances in each (in)consistency category.

|  |  | 100 Articles |  | 49 Articles |
| --- | --- | --- | --- | --- |
|  |  | Train set | Dev set | Test set |
| * | *Consistent* | 1466 | 367 | 1579 |
| Term inconsistency | (A) *Contradictory* | 160 | 40 | 128 |
|  | (B) *Over-specific* | 1172 | 294 | 1231 |
|  | (C) *Unsupportive* | 403 | 101 | 1579 |
| Evidence code inconsistency | (D) *Error-Code* | 958 | 239 | 897 |

### Modelling and Data Generation Strategy

#### Baseline

We set up two baselines using a prior-biased classifier and section information rule-based model work for each modelling task. The prior-biased classifier will make predictions according to the distribution of labels in the training set (Table 2). For example, the probability for prior-biased classifier to predict a new instance as *consistent* in the first task is 0.46, and second task is 0.60. The section information rule-based model exploits different distributions of section information among instances shown in Table 3. This model will predict any instance as *unsupportive* if its evidence section information belongs to either Background, Supporting Information, Supplementary, or Other. Otherwise, the instance will be predicted as *consistent*. However, this rule-based model will not predict any instance as *contradictory*, *over-specific*, or *erroneous code*. To overcome this issue, we apply another prior-biased classifier to process rule-based model outputs from the *consistent* type into these three types. For example, the probability of an instance that belongs to *over-specific* under the prior-biased rule-based model in the first modelling task is 0.42.

**Table 3.** Distribution of evidence section information across *Consistent* type and (C) *Unsupportive* type instances within training set.

| Section Information | *Consistent* | (C) |
| --- | --- | --- |
| Title | 1.2% | 1.8% |
| Abstract | 11.4% | 3.4% |
| Introduction | 2.7% | 13.4% |
| Background | - | 1.2% |
| Materials & Method | 3.9% | 12.6% |
| Results & Discussion | 80.6% | 53.3% |
| Conclusion | 0.2% | 0.6% |
| Supporting Information | - | 2.8% |
| Supplementary | - | 10.5% |
| Other | - | 0.6% |

#### Basic System

We model each generated instance as a paired sequence of “*[CLS]* evidence text *[SEP]* GO term + GO definition *[SEP]*” or “*[CLS]* evidence t ext *[SEP]* evidence code *[SEP]*” as input to BioPubMedBERT. We model the different types of (in)consistencies in two ways: 1) in a multi-class setting (**Model-Term**) that aims to distinguish between consistent, Type A, Type B and Type C inconsistencies by comparing evidence information with GO information; 2) and in a binary setting (**Model-Code**) for distinguishing between consistent and Type D inconsistencies by comparing evidence information with evidence code. In the basic system, two models are fine-tuned on training and development sets generated using random GO descendant selection based on the strategy mentioned in for Type B instances.

#### Optimization of Training Set

In the system with training set optimization, alternative Model-Term and Model-Code variants are fine-tuned on a collection of training and development set using the term overlap weighting strategy introduced in section for Type B instances within 20 articles. The generation of Type B instances from the remaining 80 articles follows the original strategy for preventing the model biases towards greater overlapping GO terms. The test set is retained unmodified. This optimisation aims to boost the model’s performance in distinguishing different types of instance where the semantics of GO terms is very similar, such as when “feeding behavior (GO:0007631)” and “suckling behavior (GO:0001967)” are associated with the same piece of evidence.

#### Addition of Evidence Section Information

The evidence section information is first marked by the document section title in BC4GO corpus where the evidence text located and further normalised into 10 categories. The distribution of normalised section information in *consistent* and *unsupportive* instances is illustrated in Table 3. The distribution of evidence section information is consistent with a previous statistical report on BC4GO corpus [23] where a majority of GO annotations are supported by evidence text within the Results & Discussion section. The *contradictory*, *over-specific*, and *erroneous-code* instances are generated without manipulating the evidence text from the original consistent GOA instances. Thus they retain the canonical distribution of evidence section information with Consistent GOA. The *unsupportive* instances are generated by replacing the original evidence text with unsupportive evidence sentences and therefore have different distribution of section information from consistent GOA. We concatenate the 1-dimension section encoding with 768- dimension *[CLS]* encoding in the BioPubMedBERT’s last hidden layer and forward to a linear layer in Pytorch (Fig 1).

**Figure 1.**
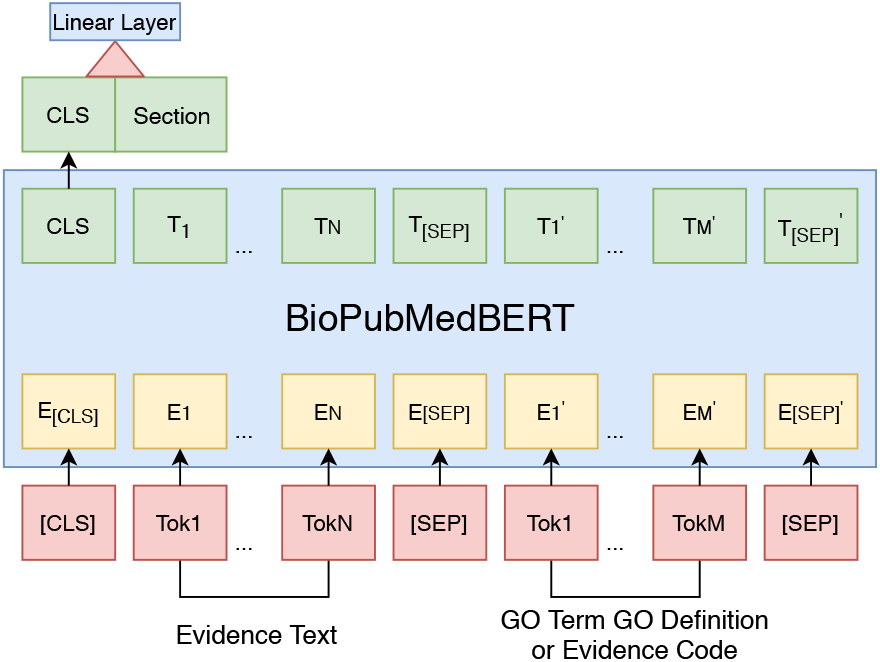
Concatenation of evidence section encoding with [CLS] encoding in classical BioPubMedBERT model. (*Tok*_*_ denotes a naturally written token, *E*_*_ and *T*_*_ denote a token embedding, *Section* denotes section encoding, *[CLS]* and *[SEP]* are special tokens that mark the boundary of a sequence pair.)

### Experiment Design

We develop models to recognise the five types of (in)consistent GOA using the baseline setting and basic system. We then run additional experiments using training set optimisation and evaluate the impact of the addition of evidence section information using *F*_1_ measure and *Precision*. We use BioPubMedBERT-uncased with 768 hidden states and the BERT-base architecture. We use the AdamW [38] optimiser with 0.01 weight decay and 300 warmup steps. We fine-tune model with 3 epochs, batch size 16 for the training set and batch size 64 for the development set. We use the huggingface AutoModelForSequenceClassification [39] framework for the fine-tuning implementation.

### Evaluation Metrics

In contrast to the existing literature-based GOA resources in GO consortium database where GO annotation is linked to evidence at the article level, we grounded annotation evidence to sentences via the BC4GO corpus. This strategy can better reflect the model’s ability to detect (in)consistent GOA. Considering an article that has two evidence sentences supporting the annotation of the same GO term, where the first evidence sentence is correctly identified as being consistent to the GO term and the second evidence sentence is not, the *Precision* for consistent GOA recognition is 1 at the article level but 0.5 at the evidence level.

We also use *Recall* and *F*_1_ as evaluation metrics for the model’s performance grounded at evidence level for each (in)consistency type. The evaluation is of the predictions over the test set, which is independent from any data point used in the previous fine-tuning stage.

To support further analysis of Model-Term’s performance, we calculate the uncertainty (*H*) of each predicted label (*i*) using Shannon entropy; see Eq. 2, in which *p_i_* denotes the probability that a GOA instance is of *i*th (in)consistency type.

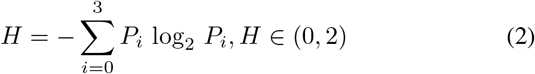

We cluster test set instances into 15 collections using an uncertainty sampling strategy derived from [33]. We use two hyper-parameters *τ*(0.2 ≤ *τ* ≤ 1.7, *step* = 0.1) and *α* = 0.1 to represent the boundary of each sample in which the uncertainty of any prediction is between *τ – α* and *τ*. This strategy can sample the same instance into more than one consecutive collection where *τ* represents the aggregated uncertainty of that collection.

We draw on the linguistic test suite of [25, 26] to define several metrics that characterise various linguistic aspects of GOA or GOA-evidence text pairs.

**GOLen:^6^** The count of tokens in a GO term split by the blank sign.
*#GOLen-2 | feeding behavior (GO:0007631)*
**AlignRatio:^6^** The count of tokens in evidence text divided by the count of tokens in GO definition text.
*#AlignRatio-0.32 Evidence: CeCDC-14 and ZEN-4 are interdependent in their localization GO definition: Any process that modulates the frequency, rate or extent of any process in which a protein is transported to, or maintained in, a specific location.*
**GEORatio:^6^** The count of word overlaps between GO term and evidence text divided by the GOLen.
*#GEORatio-0.25 GO term: regulation of protein localization Evidence: CeCDC-14 and ZEN-4 are interdependent in their localization*
**%ContainRoman:^7^** The percentage of instances that has the occurrence of Roman characters in the GO term in each sampled collection.
*#%ContainRoman-0.5 feeding behavior (GO:0007631) photosynthesis, light harvesting in photosystem <u>II</u> (GO:0009769)*
**%ContainDigit:^7^** The percentage of instances that has the occurrence of digital numbers 09 in the GO term in each sampled collection.
*#%ContainDigit-0.5 cellular response to interleukin-<u>1</u> (GO:0071347) cellular response to peptide (GO:1901653)*
**%ContainStop-OF:^7^** The percentage of instances that has the occurrence of stopword “of” in the GO term in each sampled collection^[5]^.
*#%ContainStop-OF-0.5 feeding behavior (GO:0007631) regulation <u>of</u> feeding behavior (GO:0060259)*

The Pearson correlation between the scores on these metrics and prediction uncertainty can then be investigated to provide insight into how uncertainty varies with linguistic characteristics. This analysis can be done either on a per-instance^[6]^ basis within a collection, or in aggregate across a collection.^[7]^

## Results

Table 4 shows that the model is competitive in distinguishing consistent GOA from all other types of inconsistencies compared to the baseline, with the best performance of 0.74 *Precision* for Model-Term and 0.82 *Precision* for Modelcode. The training set optimisation and the addition of evidence section information further contribute to improving the *Precision* (+0.2 & +0.15) in recognising consistent GOA. However, these two strategies do not demonstrate positive impact in distinguishing inconsistent GOA other than Contradictory (Type A). The performance of Modelcode on Type D inconsistency recognition is low, indicating that evidence code errors are difficult for the model to accurately identify.

**Table 4.**
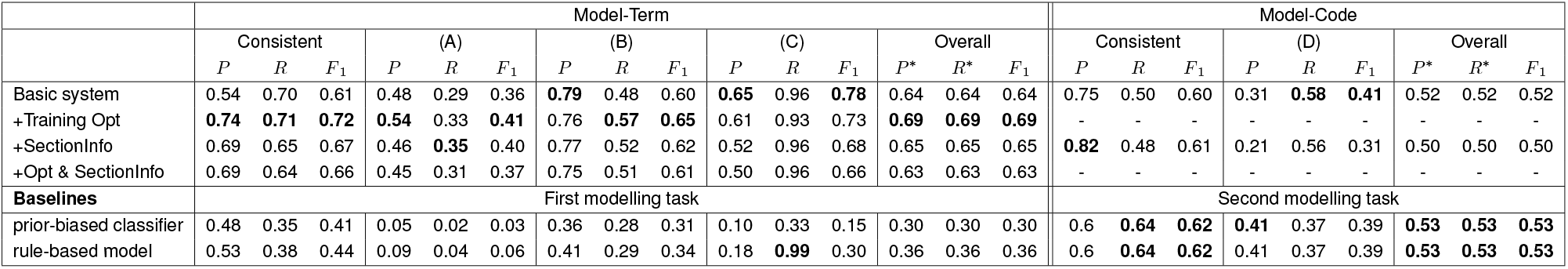
The performance of Model-Term and Model-Code in different modelling tasks, and in comparison with two baselines using *Precision* (P), *Recall* (R), *F*_1_ measures grounded at evidence level in each (in)consistency type and *Micro-Precision* (*P* *), *Micro-Recall* (*R*\*) averaged over every predicted instances in the test set. The highest metric scores for the identification of each type of (in)consistency is bolded.

Analysis of the uncertainty of predictions from Modelterm demonstrates a significant n egative c orrelation between *Precision* and the aggregated uncertainty of predictions among Consistent, Type B and Type C (Fig 2). Specifically, the model achieved above 0.9 *Precision* in recognising Type C GOA and above 0.8 *Precision* in recognising Consistent GOA among instances with prediction uncertainty lower than 0.2. The radius of scattered dots on the trending line represent the size of each sampled collection (Table 7), indicating that most of the predictions have low uncertainty. The Type A predictions do not correlate with prediction uncertainty due to the small collection size when *τ* > 0.4 is limited (less than 50 instances in each). There is an uptrend of precision for Unsupportive typed instances when *τ* range between 1.2 to 1.4. This increase may be due to either the small collection size as well (less than 40 instances in each) or the occurrence of linguistic features (illustrated in the test suite in Section) in the GO term, GO definition, or evidence text.

**Table 5.**
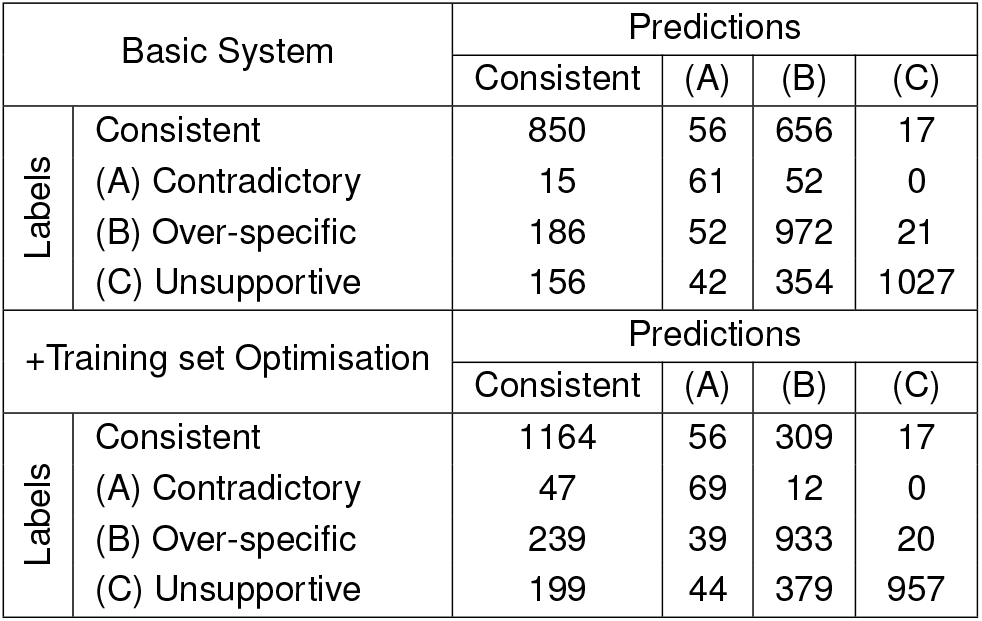
The confusion matrix of Model-Term on the test set with basic, training set optimisation fine-tuning strategy.

**Table 6.** * Pearson correlations and associated *p-value* between each score and aggregated uncertainty (the aggregated uncertainty value of each sampled collection equals to the value of hyper-parameter *τ* being set during the uncertainty sampling process, the test suite scores are averaged by taking the sum of all the instances scores and divided by the number of instances within each collection); ^ The Pearson correlation and associated *p-value* between each evaluation metric and per-instance prediction uncertainty.

|  | Predictions | Pearson R | p-value |
| --- | --- | --- | --- |
| *Precision* * | Consistent | -0.95 | 9e-08 |
|  | Type A | -0.06 | 0.82 |
|  | Type B | -0.85 | 7e-05 |
|  | Type C | -0.70 | 3e-03 |
| GOLen<sup>^</sup> | Consistent | 0.27 | 6e-28 |
|  | Type B | -0.04 | 0.14 |
|  | Type C | 0.16 | 3e-07 |
| AlignRatio<sup>^</sup> | Consistent | 0.05 | 0.03 |
|  | Type B | -0.09 | 9e-05 |
|  | Type C | 0.16 | 2e-07 |
| GEORatio<sup>^</sup> | Consistent | -0.13 | 4e-08 |
|  | Type B | -0.04 | 0.09 |
|  | Type C | 0.1 | 9e-04 |
| %ContainRoman* | Consistent | 0.79 | 5e-04 |
|  | Type B | -0.26 | 0.35 |
|  | Type C | -0.46 | 0.08 |
| %ContainDigit* | Consistent | 0.81 | 3e-04 |
|  | Type B | -0.44 | 0.10 |
|  | Type C | -0.65 | 9e-03 |
| %ContainStop-OF* | Consistent | 0.81 | 2e-04 |
|  | Type B | 0.80 | 3e-04 |
|  | Type C | 0.58 | 0.02 |

**Table 7.** The size of each sampled collection using uncertainty sampling strategy by (in)consistency type.

| $\tau$ | Size of sampled collection | | | |
| --- | --- | --- | --- | --- |
|  | Consistent | Contradictory | Over-specific | Unsupportive |
| 0.2 | 1022 | 90 | 1069 | 758 |
| 0.3 | 430 | 150 | 366 | 162 |
| 0.4 | 192 | 72 | 205 | 84 |
| 0.5 | 132 | 22 | 122 | 63 |
| 0.6 | 120 | 18 | 87 | 42 |
| 0.7 | 103 | 10 | 78 | 31 |
| 0.8 | 87 | 3 | 89 | 36 |
| 0.9 | 92 | 2 | 83 | 46 |
| 1.0 | 96 | 6 | 71 | 43 |
| 1.1 | 102 | 10 | 92 | 42 |
| 1.2 | 79 | 9 | 90 | 43 |
| 1.3 | 39 | 12 | 51 | 30 |
| 1.4 | 29 | 10 | 19 | 18 |
| 1.5 | 14 | 3 | 8 | 10 |
| 1.6 | 9 | 1 | 8 | 8 |

**Figure 2.**
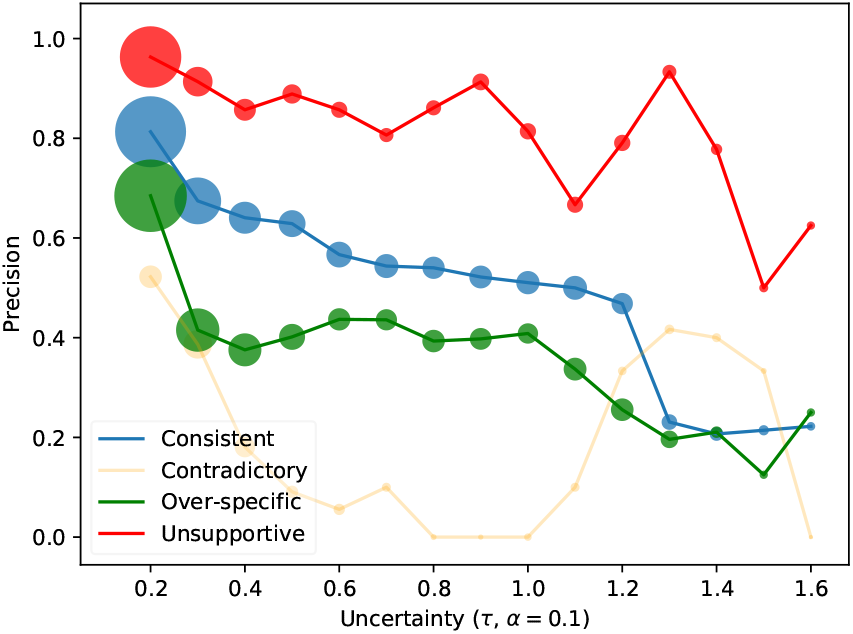
The change in *Precision* with respect to different samples under an uncertainty sampling strategy with 0.2 ≤ *τ* ≤ 1.7, *α* = 0.1 and *step* = 0.1. The radius of dots represent the size of sampled predictions that support the calculation of metric score. (The detailed Pearson correlation values can be found in Table 6 and the size of each sampled collection by (in)consistency type can be found in Table 7).

Fig 3 and Table 6 demonstrate the metrics in the test suite all have significant correlation with either model’s aggregated or per-instance prediction uncertainty. The high-lighted correlation trend lines indicate the correlation is significant (*p* < 0.05) using a Pearson correlation measure. A detailed discussion is provided in Section.

**Figure 3.**
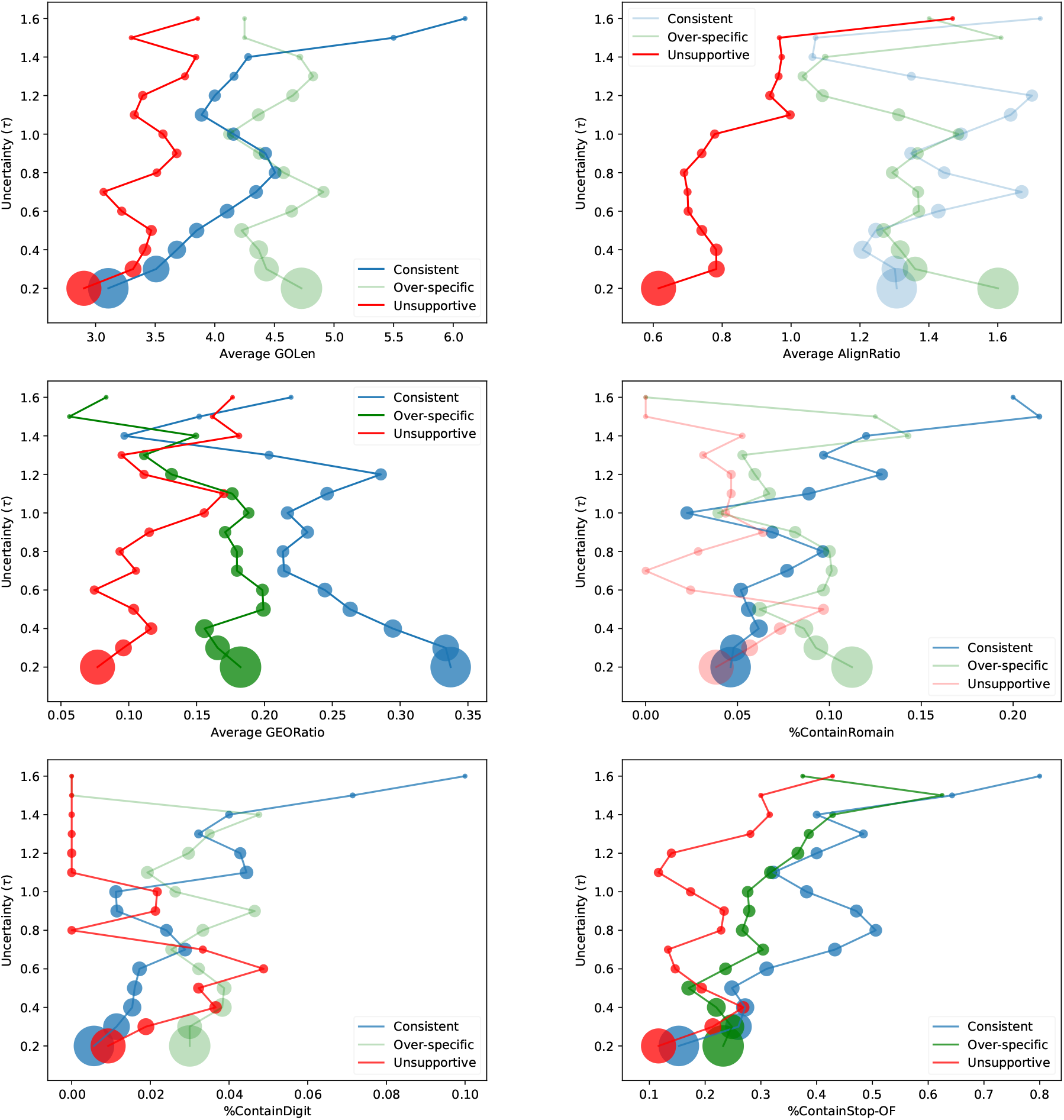
The correlation trends between each metric in the linguistic test suite and aggregated uncertainty (*τ*) of sampled collections. The highlighted correlation trend lines have a significant Pearson correlation (*p* < 0.05). (A list of Pearson values either by instance or by aggregated collection of sample can be found in Appendix, Table 6).

## Discussion

We explored the effectiveness of distinguishing *consistent* GOA and the four kinds of inconsistencies. The basic setting achieves good results in identifying Type B (overspecific) and Type C (evidence unsupportive) GOA (Table 4). However, the models failed to generalise to extreme cases where one component of the input sequence pair is highly similar to components in other instances.

For example, if two pieces of semantically similar GO information (such as “regulation of feeding behavior: Any process that modulates the rate, frequency or extent of the behavior associated with the intake of food” and “positive regulation of feeding behavior: Any process that activates or increases the frequency, rate or extent of feeding behavior”) are paired with the same piece of evidence text, the model has difficulty discriminating between them. This is reflected in the fact that most of the error cases are caused by mis-categorisation between *consistent* and *over-specific* GOA during evaluation on the test set (Table 5).

The training set optimisation contributes significant *F*1 gain in distinguishing *consistent* GOA from inconsistent GOA (Table 4). This improvement results from the correct recognition of consistent instances that were previously falsely predicted as Type B inconsistencies in the baseline setting (Table 5). However, the training set optimisation strategy does not improve the ability to distinguish different types of inconsistencies, and performance at identifying Type C inconsistencies worsened. This is because Type B and Type C inconsistencies do not strictly follow the inconsistency independence assumption (Eq 1). The Type C instances can also be seen as a Type B scenario where the associated GO term is over-specific, which makes the evidence text not supportive enough. However, the training set optimisation strategy reinforces the categorisation of such instances into either *consistent* or Type B only, which leads to the mis-categorisation of some Type C instances as Type B. A potential solution may be to group Type B and Type C inconsistencies into one category or relax the independence assumption between the two classes via a multi-label classifier.

The evidence section information is a strong indicator for discriminating Consistent GOA from other types of inconsistencies. It outperforms the basic system but can also cause biases as the mixture of *Opt&SectionInfo* underperforms the *Training Opt* method (Table 4). This is because the distribution across section segments varies between Consistent and Type C inconsistencies (Table 3). The consistent GOA only appear in the sections of Title, Abstract, Introduction, Materials & Method and Results & Conclusion, while *Unsupportive* (Type C) inconsistencies can appear anywhere in the document.

Model-Code was not successful at identifying *Erroneous Code* (Type D) inconsistencies. This is because the relationship between an evidence code and evidence text is more complex than the relationship between a GO definition and evidence text. For example, in Type A inconsistencies, there are many lexical alignments from “negative regulation” within GO terms to “decrease”, “prevent”, “deactivate” within evidence texts. In Type B instances, overspecific GO terms often have large term overlap with the correct GO term. The pairwise semantic relation patterns of GO definition and evidence text in Model-Term are restricted. However, the Model-Code input of pairwise relations between evidence code and evidence texts do not have any text alignments or term overlaps. The assessment of consistency between the two requires a comprehensive knowledge inference process which relies on both the identified evidence text and prior knowledge or other text in the article. For example, the decision of whether an evidence text such as “*Here we show that a knock-out of the ybeB gene causes a dramatic adaptation block during a shift from rich to poor media and seriously deteriorates the viability during stationary phase. YbeB of six different species binds to ribosomal protein L14. This interaction blocks the association of the two ribosomal subunits and, as a consequence, translation*” should be labelled as “IDA” [PubMed Central article PMC3400551] is decided based on information such as whether or not there is a genetic mutation or allele variation, a 1-on-1 physical interaction, or the expression pattern of gene product. It requires that the result be determined through direct assay for the function, process, or component of the gene product. This requires a sophisticated process of considering the evidence text in relation to several decision rules rather than a direct association between the text and the evidence code.

### Advanced assessment of Model-Term

Model-Term demonstrated strong potential for feasibility for real-world GO curation, particularly for the more confident predictions, as shown in Fig 2. The results of linguistic test suite analysis revealed some critical linguistic features that have significant correlation with the model’s prediction uncertainty. Specifically, the overlaps between GO term and evidence text (GEORatio) and the typical composition structure signalled by the occurrence of stopword “of” correlate well with prediction uncertainty. The correlation with this typical structure confirms that a valuable future research direction may be to develop better models of the hierarchical relationships between parent GO terms and more specific child GO terms. Additionally, we found that a longer GO term correlates with higher uncertainty in predictions of *consistent* and Type C; the differences of text length between GO definition and evidence text potentially influence the model’s uncertainty in recognising Type-C GOA; the occurrence of Roman characters and digits in the GO term demonstrate possibilities in influencing the prediction uncertainty of *consistent* and Type-C GOA as well.

We found a small number of error cases that were caused by the presence of biological or chemical formulas within GO definitions. These are not particular to any type of (in)consistency. For example, the definition for “geranyltranstransferase activity (GO:0004337)” is “*Catalysis of the reaction: geranyl diphosphate + isopentenyl diphosphate = 2-trans,6-trans-farnesyl diphosphate + diphosphate.*” We found that over 30% of formula-containing GOA instances are miscategorised by Model-Term.

### Comparison with related work

At present, not every piece of information in the generated GOA instance is exploited by our modelling: for example, GO synonyms or larger context (such as the full paragraph) from where the evidence text was extracted were not used. Some researchers have developed methods to identify hypothesis statements or new knowledge from scientific literature using language meta-knowledge [40]. According to the GO curation guidelines, evidence is unsupportive if it only express the author’s assumption of a gene function. Thus, the analysis of evidence meta-knowledge may contribute to the identification of Type-C GOA.

Our proposed method uses *vertical* consistency estimation between the GO definition and evidence text as two texts are at different levels of specificity in expressing a gene product function. The GO definition describes the gene function more abstractly while the evidence expresses more detailed information. A previous related benchmark called GOCat [10] uses *horizontal* sentence pair similarity estimation [29] where two pieces of gene function description are on the same language specificity level. It first compares the semantic similarity between new evidence text and an old GO annotated evidence text. Then it selects relevant GO terms from the old evidence text to annotate the new one. This strategy has two shortcomings: it cannot deal with new knowledge as described in Section ; and it can be biased toward frequently selected GO terms [12]. Our system can overcome these limitations and still maintain a promising performance on a test set in which over 75% of GO terms are new (Table 4). Model-Term in the basic system achieved 0.68 *micro-averaged Precision* on the test instances with new GO terms, compared to 0.74 *micro-averaged Precision* on test instances with seen GO terms. The results demonstrate that our model is effective at processing new knowledge.

## Conclusion

Continual monitoring of the consistency of GO annotation records in modern organism databases is important to maintain currency and quality of the information in these resources. We formally identify five major types of (in)consistent GO annotations that reflect the major GO annotation quality concerns for GO curation community. We propose a novel and efficient method to apply state-of-the-art text mining model to automatically detect these five major types of (in)consistent GO annotations, evaluated using an automatically generated data set. Our method satisfies the open-world assumption and is therefore robust to changes in the GO terminology.

We have demonstrated a novel method that can be adopted for real-world human-in-the-loop curation. Our implementation achieved 0.74 *Precision* for Model-Term and 0.82 *Precision* for Model-Code in distinguishing consistent from inconsistent GOA. This method can improve the efficiency of human curators by enabling curators to focus their efforts on correcting identified inconsistencies and by categorising these inconsistencies, therefore reducing the number of records that need to be manually reviewed.

Another strength is that the model has achieved competitive performance among predicted results with less prediction uncertainty, which can be used by human curators to further focus their efforts. We were able to further improve performance through training set optimisation and the addition of evidence section information. Through a detailed performance analysis using a linguistic test suite we identified superficial linguistic features that may impact the model’s prediction uncertainty.

In future work, we aim to produce a more comprehensive evidence-based GO annotation corpus focusing on inconsistencies, seek assistance from expert curators to test and extend the proposed methods on real-world database records with broader gene function perspective, and will specifically seek to improve the identification of evidence code inconsistencies. We will also examine the use of meta-knowledge analysis to improve the model’s performance in identification of instances that lack supportive evidence. We will refine the modelling of semantic hierarchical relationship between parent and children GO terms.

## Authors’ contributions

JC contributed to the exploration, experiment, analysis and drafted the manuscript. NG JZ and KV contributed to the conceptualization of the research and in revising the manuscript. All authors read and approved the final manuscript.

## Availability of data and materials

All data presented and analyzed in the present study was generated from the BC4GO corpus and retrieved GO Consortium Database. A demonstrable code for the generation of adversarial instances is accessible at https://github.com/jiyuc/BioConsistency. There are no additional materials created associated with this study.

## Ethics approval and consent to participate

Not applicable

## Acknowledgements

Not applicable

## Consent for publication

Not applicable

## Footnotes

[1] http://geneontology.org/docs/guide-go-evidence-codes/

[2] http://geneontology.org/stats.html

[3] http://geneontology.org/docs/ontology-relations

[4] ftp://ftp.geneontology.org/go/www/images/diagevCodeFlowChart.pdf

[5] Note that GO terms are drawn from strictly controlled vocabularies where the stopword “of” expresses a compositional structure, with a parent GO term often appearing within a child term. Thus, this trick of counting the occurrence of such typical compositional structure via the stopword “of” is only feasible for GO terms but not GO definitions or naturally written evidence text.

[6] The test suite metric scores can be calculated at the instance level or aggregated by taking the average: taking the sum of all the instances metric scores and divided by the number of instances within a collection

[7] The aggregated uncertainty value of each sampled collection equals to the value of hyper-parameter *τ* being set during the uncertainty sampling process, the test suite metric scores can only be calculated on the percollection basis

